# Inflammatory cytokines rewire the proinsulin interaction network in human islets

**DOI:** 10.1101/2022.04.07.487532

**Authors:** Duc Tran, Anita Pottekat, Kouta Lee, Megha Raghunathan, Salvatore Loguercio, Saiful Mir, Adrienne W. Paton, James C. Paton, Peter Arvan, Randal J. Kaufman, Pamela Itkin-Ansari

## Abstract

Aberrant biosynthesis and secretion of the insulin precursor proinsulin occurs in both Type I and Type II diabetes (T1D, T2D). Inflammatory cytokines are implicated in pancreatic islet stress in both forms of diabetes but the mechanisms remain unclear. Here we examined how the diabetes associated cytokines interleukin-1β and interferon-γ alter proinsulin interactions with proteins that regulate its folding, trafficking, and secretion. Human islets treated with cytokines exhibited secretion of proinsulin, IL6 and nitrite, as well as evidence of endoplasmic reticulum (ER) stress. Unbiased proinsulin Affinity Purification-Mass Spectrometry revealed a proinsulin interactome reshaped by cytokines relative to controls. Cytokine treatment increased proinsulin binding to multiple ER chaperones and oxidoreductases, including the major ER chaperone BiP. Moreover, increased BiP binding was an adaptive response required to maintain proinsulin folding in the inflammatory environment. Cytokines also regulated novel interactions between proinsulin and T1D and T2D GWAS candidate proteins not previously known to interact with proinsulin (*e*.*g*., Ataxin-2) and these GWAS proteins formed a tight network with each other. Finally, cytokines induced proinsulin interactions with a cluster of microtubule motor proteins. Consistent with a role for these proteins in proinsulin trafficking and release, chemical destabilization of microtubules with Nocodazole exacerbated cytokine induced proinsulin secretion. Together, the data quantitatively map the proinsulin interactome rewired by cytokines, shedding new light on how human proinsulin biosynthesis is dysregulated by an inflammatory environment.

## Introduction

All forms of diabetes ultimately result from β-cell failure. Type 1 diabetes (T1D) is a chronic inflammatory disease with autoimmune β-cell destruction, and it is now accepted that T2D is also characterized by inflammation with deleterious effects on the β-cell (1-7). Given that more than 400 million people world-wide suffer from diabetes, it is critical to fully understand the physiological consequences of inflammation on the β-cell.

Insulin is initially synthesized as preproinsulin that enters the endoplasmic reticulum (ER) where cleavage of its signal peptide converts it to proinsulin. The ER guides oxidative folding of three critical disulfide bonds in proinsulin that are facilitated by chaperones and oxidoreductases, *e*.*g*., BiP and PDIA1. We and others identified several ER chaperones and oxidoreductases required for proper proinsulin folding (8-11). Only properly folded proinsulin transits through the ER-Golgi intermediate compartment (ERGIC) to the Golgi apparatus from which immature secretory granules are formed. Inside the immature granules, processing enzymes cleave the C-peptide from proinsulin to generate mature insulin that is secreted in response to glucose. Under healthy conditions proinsulin is efficiently converted to insulin and little proinsulin is released into the serum.

Three highly conserved pathways regulate ER homeostasis through the unfolded protein response (UPR): PERK, IRE1 and ATF6. Activated PERK reduces ER burden by transiently attenuating general mRNA translation, except for proteins that resolve ER stress. Loss of PERK *in vivo* causes β-cell failure and permanent neonatal diabetes, and we have shown that even short-term PERK inhibition in isolated human islets induces proinsulin misfolding (11). IRE1 activation leads to splicing of the transcription factor XBP1 that induces genes encoding ER protein translocation, ER chaperones and ER-Associated Degradation (ERAD) effectors. ATF6 processing and activation leads to its nuclear translocation to induce expression of genes encoding functions complementary, as well as overlapping, with spliced XBP1. Under homeostatic conditions, ER stress signaling through the three UPR sensors is limited by their binding to BiP as BiP interaction with PERK, IRE1 and ATF6 represses their signaling. When BiP encounters misfolded protein however, it releases from PERK, IRE1 and ATF6, activating ER stress signaling (12). Thus, increased protein binding to BiP is considered a gold standard as an indicator of increased unfolded/misfolded protein.

β-cells produce up to 6000 molecules of proinsulin per second (13) making them susceptible to ER stress and proinsulin misfolding under conditions of increased demand for proinsulin, as occurs when a portion of β-cells have been destroyed, or in conditions of insulin resistance. There is evidence of ER stress in islets from patients with T1D and T2D (14) and ER stress precedes T1D in the nonobese diabetic (NOD) mouse islets (4, 15-18). We showed that high glucose alone (as occurs in both forms of diabetes) is sufficient to induce high molecular weight aggregates of misfolded proinsulin in normal human islets. We also observed proinsulin misfolding in islets from patients with T2D (11) and in islets from mice fed a high fat diet (HFD) (19). Strikingly, proinsulin misfolding preceded hyperglycemia in the db/db model of T2D (19). Together, the data suggest a connection between inflammation, ER stress and proinsulin misfolding.

A second common feature of islet pathology in T1D and T2D is secretion of unprocessed proinsulin (20). In fact, detectable serum proinsulin persists in a majority of individuals with longstanding T1D, even those that no longer have detectable serum C-peptide (21). Moreover, proinsulin in the bloodstream is a predictor of T2D, obesity, and cardiovascular disease and the ratio of secreted proinsulin to insulin is a measure of β-cell dysfunction associated with T2D (22). Notably however, circulating proinsulin also provides clear evidence of β-cell survival in patients with T1D and T2D. Therefore, deciphering new strategies to restore optimal performance in these β-cells is critically important.

To gain insight into the effects of an inflammatory environment on β-cell function, transcriptomic, proteomic, and epigenetic analyses of human islets treated with diabetes associated cytokines interleukin-1β (IL-1β) and interferon-γ (IFN-γ) have cataloged genes/proteins regulated by cytokines (2, 23). While these data illuminate the composition of islets, they do not identify the proteins that interact with proinsulin to coordinate its folding, trafficking, and processing. We recently used proinsulin affinity purification-mass spec (AP-MS) to comprehensively define the proinsulin interactome in normal human islets (11). Here human islets treated with IL-1β and IFNγ were similarly subjected to proinsulin AP-MS for comparative analysis with the reference interactome.

The AP-MS data revealed that cytokine treatment modified proinsulin binding to multiple ER chaperones and oxidoreductases while increasing a propensity to proinsulin misfolding. Novel proinsulin interactions with proteins previously identified as diabetes GWAS candidates were also regulated by cytokines. Finally, cytokines were found to enhance proinsulin interaction with a protein network comprised of several microtubule-regulatory proteins. We show that microtubule dynamics play a role in aberrant proinsulin release, as functional destabilization of microtubules by Nocodazole enhanced cytokine induced proinsulin release.

## Results

### Cytokines promote inflammatory signaling and increase both insulin and proinsulin release from human islets

To investigate the effects of inflammation on β-cell function, human islets were treated with the diabetes associated cytokines IL-1β (50U/ml) and IFN-γ (1000U/ml) for 48 hours. These conditions were previously shown to emulate many aspects of diabetes associated inflammation without inducing significant cell death; (24-28) and consistent with this, cytokine treated islets appeared normal size and morphology (**Fig.1A**).

To measure inflammatory signaling induced by cytokines, we used a combination of RNA-Seq, RT-qPCR, Western blots, the Griess assay for nitrite release, and ELISAs. Analysis of RNA-Seq by Ingenuity Pathway Analysis (IPA) revealed that ‘Antigen presentation’ and ‘Type I Diabetes signaling’ were among the top four pathways induced by the cytokines versus untreated islets (**Fig.1B**). Strikingly, analysis of upstream signaling effectors predicted to generate the observed RNA-Seq profiles correctly identified IL-1β, and IFNγ (**Fig.1C**). We next compared our RNA-Seq data to a recent proteomic study of human islets treated with the same cytokine cocktail, identifying a strong correlation between the RNA and protein profiles (Pearson correlation: 0.7736988, p-value < 2.2 × 10^−16^)(**Fig. 1D**) (23). RNA-Seq and RT-qPCR revealed that cytokines induced inflammatory effectors, *e*.*g*., the chemokine CXCL10 (29), and inducible Nitric Oxide Synthase (iNOS from the NOS2 gene) (**Fig. 1D**), suggesting that cytokines induced the release of inflammatory factors from islets. Indeed, the Griess assay revealed that cytokines triggered nitrite release in all islet preparations tested (*p* = 7×10^−4^) (**Fig. 1F**) and induced a 32-fold increase in secreted IL6, as measured by an ELISA, *p* <0.001 (**Fig. 1G**).

**Figure 1.**
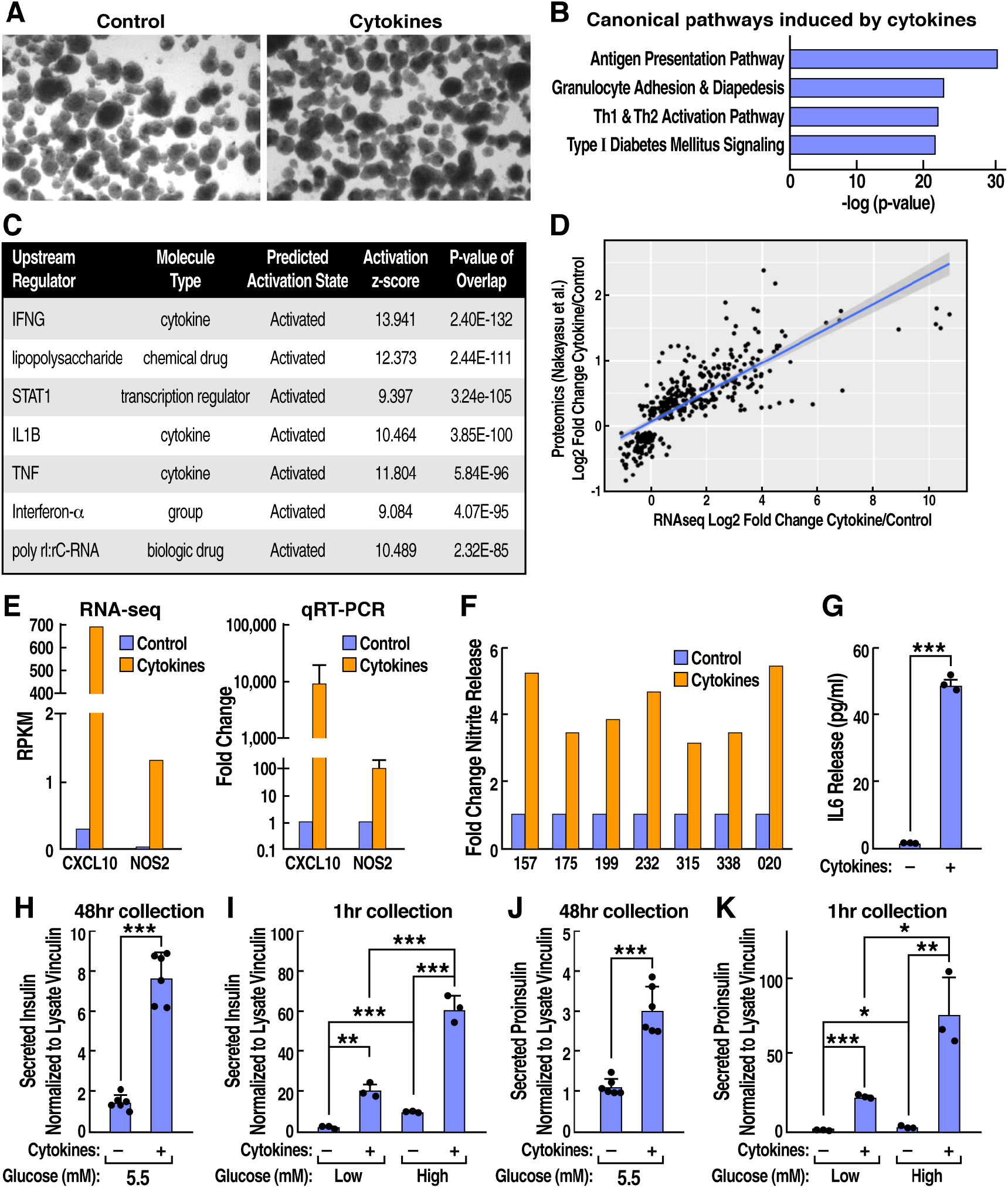
Cytokine treated human islets exhibit upregulated chemokine expression with release of IL6 and NO, altered GSIS, and aberrant proinsulin release. Cytokine treated and untreated islets were assayed for: A) morphology by light microscopy; B) IPA Canonical pathways induced by cytokines in RNA-Seq data; C) upstream pathways predicted by IPA to produce the observed cytokine regulated expression profile in RNA-Seq data; D) Comparison of cytokine versus untreated human islets in our RNA-Seq profile versus published whole proteome data (23); E) RNA-Seq and qRT-PCR analysis of CXCL10 and NOS2 (iNOS) expression; F) Nitrite release by Griess assay, G-K) ELISA data for secreted human: G) IL6; H,I) insulin and J,K) proinsulin. Data represent the mean of three-six independent biological replicates analyzed by unpaired t-test. Error bars are -/+ SEM, *p≤0.05, **p≤0.01, ***p≤0.001.

We next examined the effect of cytokines on insulin and proinsulin release from human islets. Media collected after 48-hour incubation +/- IL-1β and IFNγ treatment (in 5.5mM glucose) and assayed for insulin by ELISA (p<0.001) showed a marked increase in insulin release due to cytokines (**Fig. 1H**). To measure glucose stimulated insulin secretion (GSIS) the islet samples previously treated +/- cytokines for 48 hours underwent cytokine washout followed by sequential one-hour incubations in cytokine free 2.2 mM (low) and then 22mM (high) glucose media. Cytokines increased insulin release at both low and high and glucose (**Fig. 1I**). To determine whether the cytokine treated islets exhibited aberrant proinsulin release associated with β-cell dysfunction in diabetes, the media samples collected above were also assayed for proinsulin by ELISA. In islets treated with cytokines for 48 hours, a nearly three-fold increase in proinsulin release was observed (**Fig. 1J**) (**Supp 1**). Similar to insulin, cytokine treated islets exhibited higher levels of proinsulin secretion than controls in normoglycemia, low and high glucose incubations without significantly altered proinsulin in lysates. ELISA values were normalized to vinculin from corresponding islet lysates. (**Fig. 1K)**. The trends in insulin release were similar to those observed for proinsulin. To ensure that insulin and proinsulin in the media emanated from intact islets, media was analyzed by Western blot for two of the ten most highly induced cytoplasmic proteins by cytokines, GBP5 and IDO1. The two proteins were only detected in cell lysates and not in supernatants, arguing against cell lysis as a mechanism for proinsulin release (**Supp Fig. 1**).

### Defining the proinsulin interactome in cytokine treated islets

We recently reported the proinsulin interactome in normal human islets that was highly conserved across a diverse donor group that included both sexes and spanned three ethnicities (11). To examine the effects of cytokines on the same islet samples, equal portions of the six islet preparations were treated with cytokines. The islets were then lysed and immunoprecipitated with beads conjugated with a mouse mAb specific for human proinsulin, or with control mouse IgG–conjugated beads. A small amount of each sample was used to establish that total proinsulin recovery was similar in cytokine-treated and untreated samples (**Supp. Fig. 2A**). The bulk of each sample was then subjected to on-bead denaturation, reduction, trypsin digestion and LC-MS/MS analyses. LC-MS/MS runs were conducted concurrently with the corresponding untreated islet samples (data for the latter is published) (11). MS/MS spectra were searched against the human Uni-Prot database and analyzed with MaxQuant.

Comparison of MS/MS counts for proinsulin (bait) across the six islet samples confirmed the remarkable consistency in proinsulin recovery with and without cytokines. Three of the islet samples were also assayed in duplicate, to ensure technical reproducibility (**Supp. Fig. 2B**). Similar numbers of proteins were identified in all MS runs (**Supp. Fig. 2C**). As in our previous study, Loess-R was used for normalization and MSstats was used to calculate a statistical confidence score (*p-*value) for each protein based on intensity and reproducibility of detection across samples.

To identify the most highly conserved cytokine induced alterations in the proinsulin biosynthetic pathway, data were analyzed using the four criteria we described for normal islets (11): *i*) total intensity in Proinsulin IP vs IgG IP ≥ 2, *ii*) p ≤ 0.05, *iii*) MS/MS ≥ 10, and *iv*) gene expression in single cell RNA-Seq of human beta cells >1 per million reads (30). This stringent analysis produced a dataset of 527 identified proteins, 162 enriched cytokine treatment and 365 shared with those we reported in untreated islets (11). For visualization of the data, each protein was assigned to a compartment or function based on Genecards, Uniprot or literature (**Fig. 2**). For simplicity, proteins assigned a primarily ribosomal, nuclear, mitochondrial cellular function were not included in **Fig. 2**. A selection of identified proteins were validated by Proinsulin AP-Western blot (**Supp. Fig. 3**).

As we noted in the proinsulin interactome in untreated human islets, MS results for cytokine treated islets displayed striking sensitivity and specificity. The sensitivity is exemplified by identification of the processing enzyme PCSK1, despite the fact that it interacts only transiently with proinsulin. In contrast, specificity was demonstrated by lack of non-specific proinsulin interaction with amylin that is expressed at high levels and co-secreted with insulin in β-cells. Nor did proinsulin interact with chemokines that were among the proteins most highly induced by cytokines (23).

### Cytokines alter proinsulin folding and interactions with ER protein folding machinery

IPA of all proinsulin interactions identified ‘Protein folding in ER’ as a pathway significantly altered by cytokines (**Fig. 3A**). Among ER folding factors, the chaperone BiP was the most significant proinsulin interactor (Proinsulin IP/IgG IP) under cytokine conditions (*p*=0) (**Fig. 2**) and also exhibited the most significant *p*-value for the *increase* in binding in cytokine treated relative to control islets p=<0.001, validated in **Supp Fig. 3**. We previously showed that BiP plays a critical role in proinsulin folding in human and murine islets (10, 11, 19), suggesting that increased BiP might be required to maintain proinsulin folding in an inflammatory environment. To test this, islets +/- cytokines were treated with the Shiga-toxigenic *Escherichia coli* virulence factor subtilase toxin SubAB that cleaves and inactivates BiP. As a control, islets were treated with enzymatically inactive mutant toxin SubA_A272_B (mSubAB) (31). **Fig. 3B** shows that cytokine treatment exacerbated SubAB induced proinsulin misfolding. We previously reported PERK inhibition, which stimulates proinsulin synthesis (32), also triggers proinsulin misfolding in human islets (11). Therefore, we examined the effect of cytokines on islets treated with the PERK inhibitor GSK157. Islets incubated with GSK157 in the presence of cytokines exhibited increased proinsulin misfolding compared to GSK157 treatment alone (**Fig. 3B**). Together, the data suggest that islets in an inflammatory environment are in a precarious position with regard to proinsulin folding. This may result from a combination of both gained and lost interactions, *e*.*g*., in addition to BiP, cytokines increased proinsulin binding to the oxidoreductase PDIA1 (P4HB, **Figs. 2**,**3A**) that we recently showed is required for efficient proinsulin maturation and β-cell health in response to stress *in vivo* (diet induced obesity) (10). In contrast however, PRDX-4 that we showed was required for protein folding in oxidant stressed β-cells (as with cytokine treatment), exhibited lower proinsulin binding in cytokine treated islets (**Fig.2**). Taken together, the data are consistent with a response to cytokines that is both adaptive and maladaptive.

**Figure 2.**
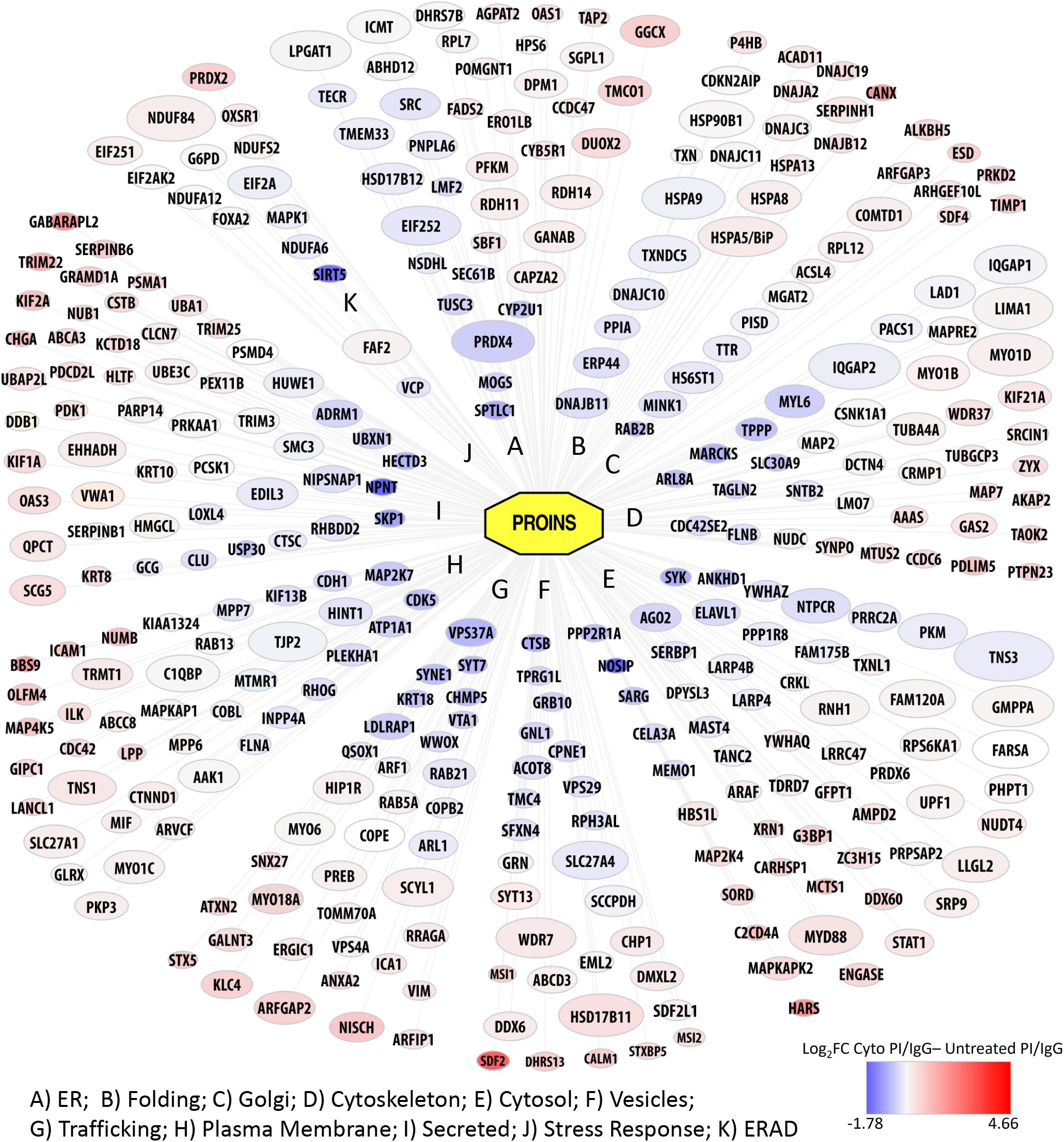
Defining the proinsulin interaction network regulated by cytokines. AP-MS identified proinsulin interactions in cytokine treated islets were compared with recently published proinsulin interactome in untreated islets (11). Icon size reflects significance of interaction with PI vs control IgG, increasing icon size =increasing significance (smallest icons *p*=0.05). Deepening red reflects PI interaction increased by cytokines and deepening blue reflects PI interactions decreased by cytokines, palest for each color=2-fold difference. Protein categories were defined by the Human Protein Atlas. For simplicity, PI interacting proteins primarily in nucleus, mitochondria, ribosome, or undetermined categories are not visualized in this figure but are included in uploaded dataset.

**Fig. 3.**
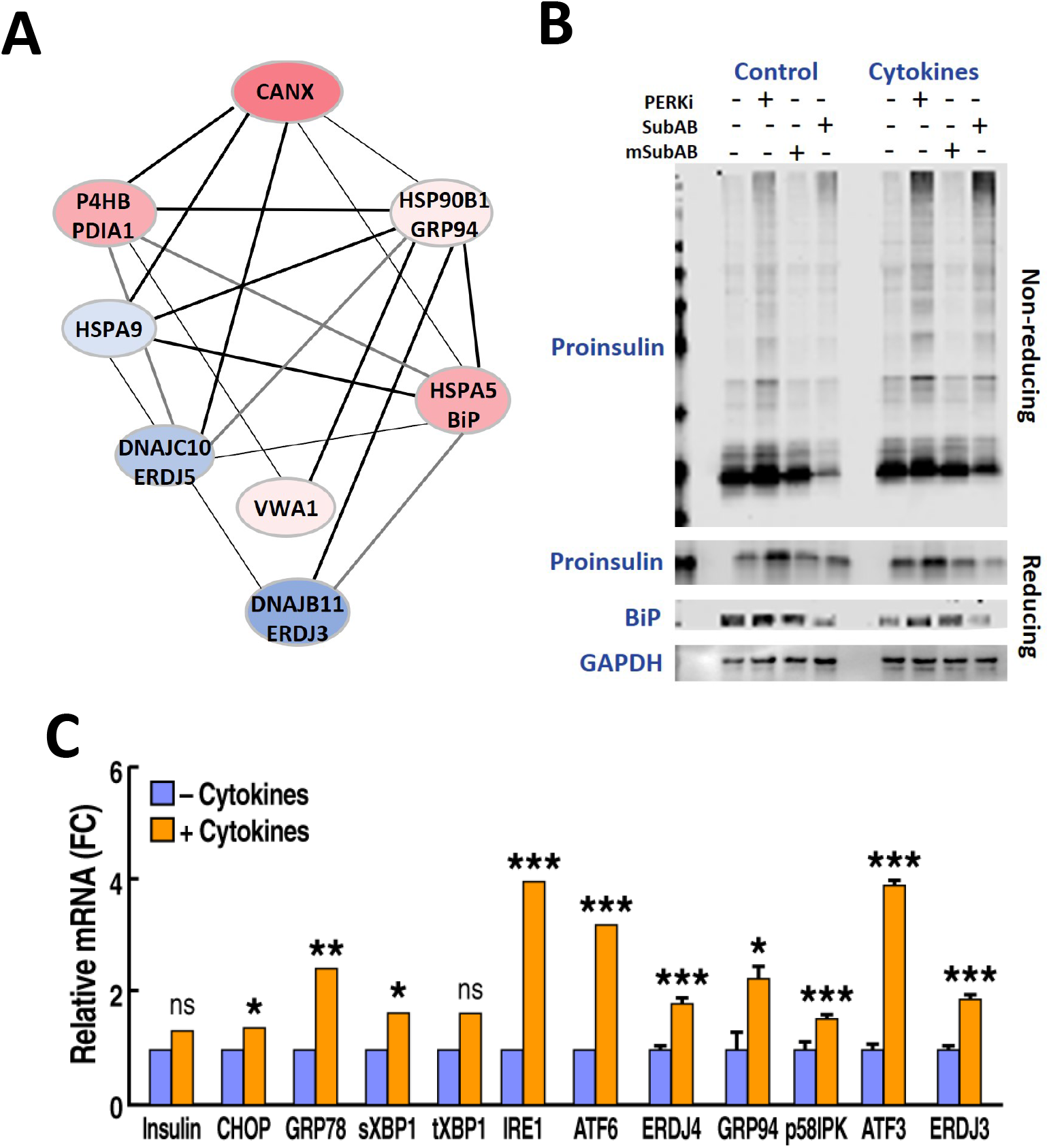
Cytokines alter human proinsulin interactions with ER factors and induce sensitivity to proinsulin misfolding. A) String Mode Cluster Analysis identifies ‘Protein folding in ER’ as altered by cytokines. B) Human islets treated +/- cytokines, +/- PERK inhibitor and +/- SubAB (that cleaves BiP) analyzed by Western blot for proinsulin on non-reducing SDS-PAGE. Western blots for proinsulin, BiP and GAPDH shown below were analyzed by reducing SDS-page. C) RT-qPCR of UPR genes in cytokine treated vs untreated human islets Error bars are -/+ SEM, *p≤0.05, **p≤0.01, ***p≤0.001.

We next examined the expression of ER stress response genes by RT-qPCR. Cytokines modestly increased the levels of the UPR genes CHOP, ATF6 and IRE1 and spliced XBP1 among others (**Fig. 3C**). ER stress appeared to be sub-maximal as we did not observe phosphorylation of the translation initiation factor eIF2α at Ser51, an event that leads to translation attenuation upon PERK activation (**Supp Fig. 4**).

### Diabetes GWAS candidate proteins interact with proinsulin and with each other

We observed that cytokines regulated proinsulin interactions with several GWAS candidates associated with T1D and T2D (**Fig. 4**). Among T1D candidates, cytokines increased proinsulin interactions with TAP2 and PRKD2, while interactions with COBL (33) and TUSC3 were decreased. Among T2D candidates, cytokine exposure induced proinsulin interactions with SDF2L1, SCYL1, IGF2BP2 and UBE3C and reduced interactions with SYNE1 (34, 35). Two GWAS candidates implicated in both T1D and T2D are ATXN2, that exhibited upregulated proinsulin interactions in cytokine conditions, and PLEKHA1, that exhibited proinsulin interactions downregulated by cytokines (type2diabetesgenetics.org) (36, 37). To investigate whether proinsulin binding GWAS proteins formed a network, we used unsupervised analysis of published protein:protein interactions curated in BioGrid (BioGrid 4.4), for each candidate. Strikingly, the GWAS factors formed a tight matrix of interactions with each other and with a small number of additional proinsulin interactors we identified by AP-MS (**Fig.4**). Taken together, the proinsulin AP-MS data suggest β-cell roles for diabetes GWAS candidates for which little has been known.

**Figure 4.**
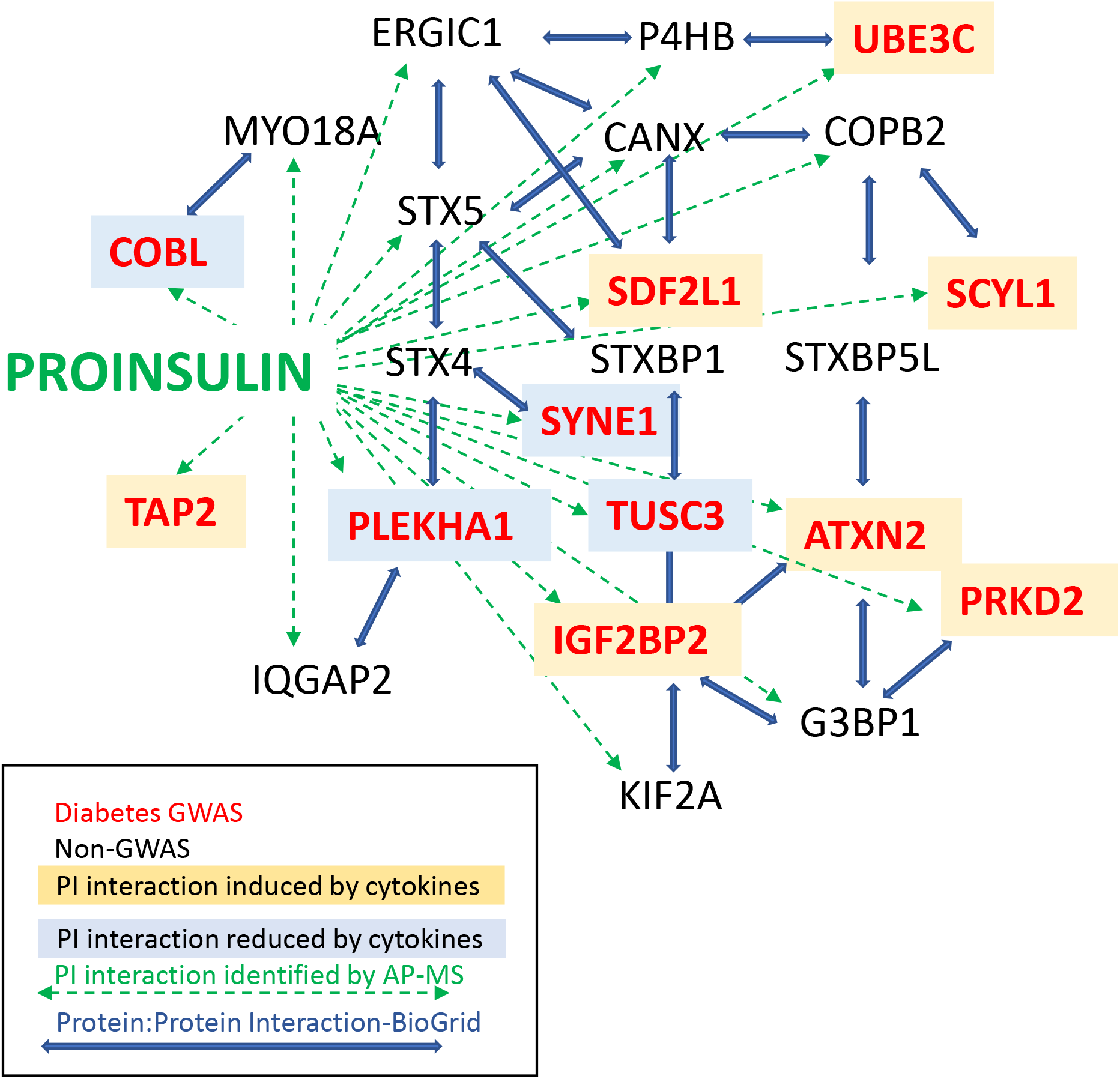
Cytokines regulate proinsulin interactions with diabetes GWAS candidate proteins. Green dashed arrows show proinsulin interactors identified by AP-MS. Of the connections shown here, STX4, STXBP1 and STXBPL5 are the only proteins that were not identified as proinsulin interactors. GWAS proteins are distinguished by red lettering. Yellow shading=increased proinsulin binding with cytokines, blue shading=decreased proinsulin binding with cytokines. Blue arrows represent protein:protein interactions identified in BioGrid.

The proinsulin interactome data also provided clues to islet autoimmunity that contributes to T1D pathogenesis. Proinsulin itself is an autoantigen in T1D (38), and we found that cytokines induced proinsulin interactions with the additional autoantigens ICA69 and chromogranin A (39).

### Cytokines induced proinsulin interactions with ER-Golgi transport and microtubule associated proteins

Ingenuity Pathway analysis revealed that the proinsulin interaction network includes a cluster of Golgi-ER trafficking proteins regulated by cytokines. Among these, cytokines significantly induced proinsulin interactions with several kinesins, a family of microtubule motor proteins, *i*.*e*., KIF1A, KIF21A, and most significantly, KIF2A (FC Cytokine PI/IgG= 4.2, *p*=1.1×10^−6^) (**Fig. 5A**). KIF2A binding to proinsulin was verified by proinsulin AP-Western blot (**Fig. 5B**). Increased KIF2A localization with proinsulin in cytokine treated human islets was also validated by immunohistochemistry (**Fig. 5C**). The immunohistochemistry showed that KIF2A protein expression in β-cells was heterogeneous and analysis of previously published single cell RNA-Seq data also showed KIF2A heterogeneity at the level of transcription (**Supp Fig. 5**) (30). KIF2A has plus-end microtubule depolymerizing activity. Because microtubule depolarization was shown to increase insulin release during GSIS (40, 41), we investigated whether microtubule dynamics play a role in cytokine induced proinsulin release. Human islets incubated in 5.5 mM glucose +/- cytokines for 48 hours were treated +/- the microtubule destabilizing drug Nocodazole for the final 24 hours. At the 48-hour point, media was collected (48-hour samples) and islets were washed for one-hour in 2.2 mM glucose prior to a one-hour incubation in 2.2 mM (low glucose), followed by 22 mM (high glucose) media, with a sample collected from the latter at ten minutes to gauge first phase release (high glucose, ten minutes), and the remainder collected at one-hour (high glucose, one-hour). At all glucose concentrations, Nocodazole alone stimulated proinsulin release, as did cytokines alone. Moreover, Nocodazole acted in concert with cytokines to induce proinsulin secretion as well as insulin secretion (**Fig. 5D**).

**Figure 5.**
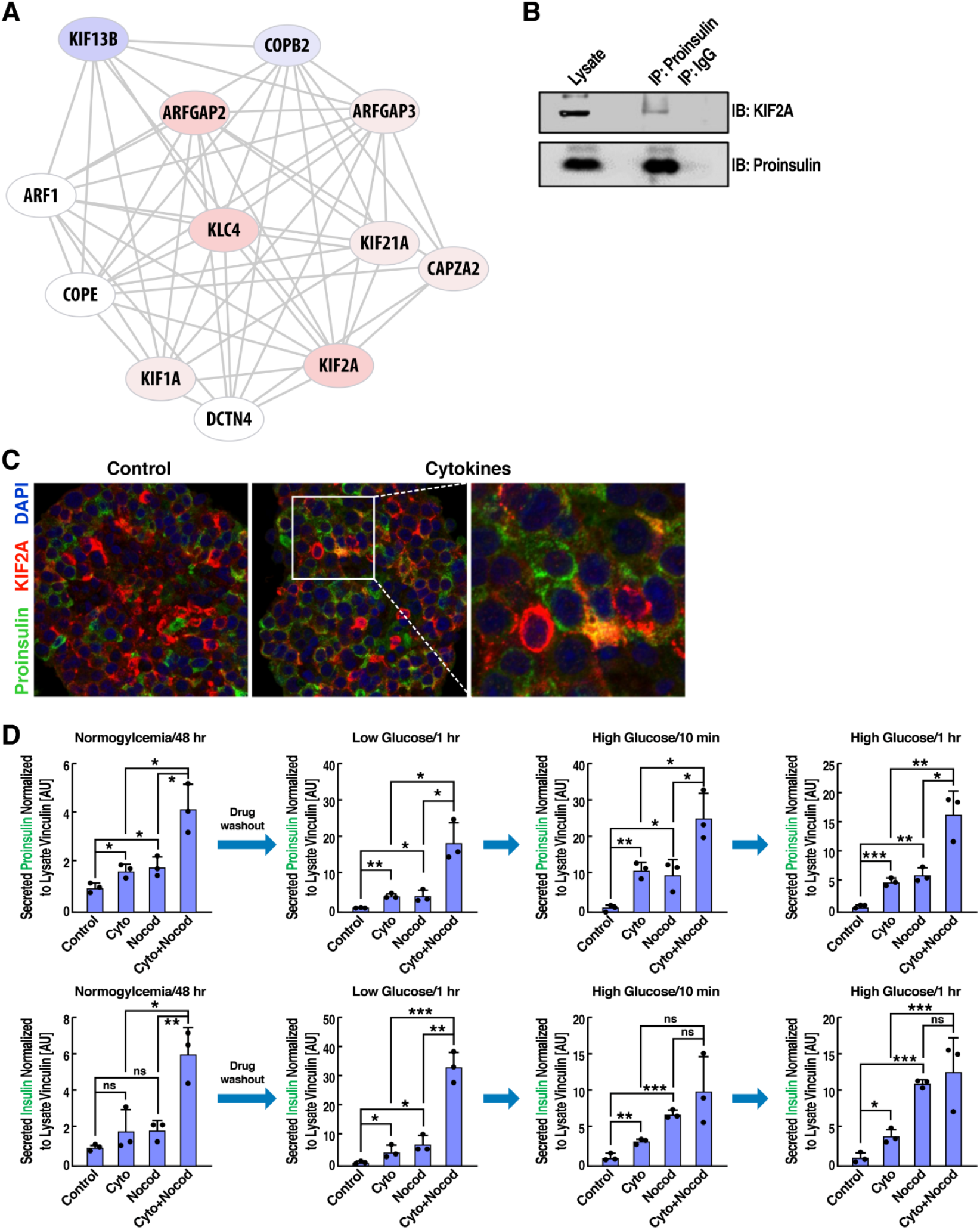
Cytokines induce proinsulin interactions with microtubule associated trafficking proteins, and microtubule destabilization promotes proinsulin secretion. A) Cytokine induced interactome changes in proinsulin Golgi-ER transport network identified by Ingenuity Pathway Analysis. B) Validation of proinsulin binding to KIF2A in human islets by proinsulin IP and Western blot. C) Immunohistochemistry of paraffin embedded human islets +/- cytokine treatment, immunostained for KIF2A and Proinsulin. D) Media was harvested from human islets and assayed for Proinsulin and Insulin by ELISA following 48-hour incubation +/- cytokines in 5.5 mM glucose, then incubations for 1 hour in 2.2mM glucose, 10min in 22mM glucose, and 1 hour in 22mM glucose. At the conclusion of all incubations, islet samples were lysed and lysates analyzed by Western blot for vinculin that was used for normalization. Data represent the mean of three independent biological replicates analyzed by unpaired t-test. Error bars are -/+ SEM, *p≤0.05, **p≤0.01, ***p≤0.001.

## Discussion

Recently we identified the proinsulin biosynthesis network in normal human islets (11). The data set the stage for comparative analyses with interactomes from islets under perturbed conditions. Here we sought to understand how inflammatory cytokines associated with T1D and T2D progression affect the network of human proinsulin protein interactions along the secretory pathway.

Cytokines altered proinsulin interactions with numerous ER resident proteins, in what appears to be a mix of adaptive and maladaptive changes. For example, significantly increased proinsulin binding to the major ER chaperone BiP was observed in cytokine treated islets. Inactivation of BiP with the toxin SubAB provoked more proinsulin misfolding in cytokine conditions than in normal human islets, suggesting that increased BiP binding is required to maintain proinsulin folding in cytokine treated cells. In contrast, cytokine treatment lowered proinsulin interactions with PRDX4, a peroxiredoxin that we recently showed promotes proper proinsulin folding, particularly in stress conditions. PRDX4, when overexpressed, was also shown to protect β-cells against streptozotocin induced injury insulitis (42). In summary, proinsulin folding appears to be maintained in cytokine conditions through recruitment of additional chaperones and oxidoreductases, yet reduced interactions with PRDX4 and co-chaperones (*e*.*g*., ERDJ3 and ERDJ5) may explain the increased sensitivity to misfolding.

We noted modest ER stress in cytokine treated islets. The relationship between ER stress and protein misfolding may result in a vicious cycle leading to β-cell failure. It is well known that ER stress, *e*.*g*., induced by Thapsigargin, a non-competitive inhibitor of the sarco/endoplasmic reticulum Ca^2+^ ATPase (SERCA), or inhibition of the UPR sensor PERK, results in proinsulin misfolding. Interestingly however, a lesson from patients with Mutant Ins-gene-induced Diabetes of Youth (MIDY) is that proinsulin misfolding can also be the primary event that drives ER stress. MIDY is characterized by a misfolding-inducing mutation in only one allele encoding insulin. The mutant proinsulin is unable to fold and entraps the non-mutant proinsulin in high MW complexes that cannot leave the ER, and ER stress ensues. Thus, the data suggest that regardless of the initiating event, ER stress and proinsulin misfolding generate a propagative cycle leading to β-cell failure.

ER and Golgi localized proteins were among GWAS candidates identified as proinsulin binding. In the ER, The T2D candidate SDF2L1 is involved in ERAD (43), the T1D candidate TUSC3, is an ER glycosyltransferase (44, 45), and the T1D candidate TAP2 (**Fig. 2 and 4**). Proinsulin binding GWAS factors localized to the Golgi included SCYL1 (46) and the T1D candidate PRKD2, a trans-Golgi network (TGN) protein required for the release of chromogranin-A (CHGA)-containing secretory granules (33, 47). Moreover, BioGrid analysis of the GWAS candidates’ protein:protein interactions yielded a tight network of proteins that comprise the GWAS factors and a small number of additional proinsulin interactors. The data therefore suggest that the GWAS candidates sit at a nexus of a highly conserved matrix of proteins involved in proinsulin biosynthesis. To the best of our knowledge, other than TAP2 and SDF2L1, these GWAS candidates have not previously been reported to interact with proinsulin.

Cytokine treatment induced proinsulin release into the media (48) mimicking the pathological proinsulin release observed in sera from patients with both T1D and T2D. The proinsulin release occurred in the context of increased proinsulin binding to the microtubule-associated kinesin motor proteins KIF1A and KIF21A, KIF2A. KIF2A, a microtubule depolymerase, was also identified in a proinsulin interactome study we conducted in MIN6 cells, which employed a different antibody to immunoprecipitate proinsulin (49). The association of insulin granules with microtubules has been known for some time (50), and Golgi-derived microtubules are critical for replenishment of the insulin granule pool and proper insulin secretion (40, 41). Given the increased binding of proinsulin with KIF proteins, we tested whether microtubules played an essential role in cytokine induced proinsulin. Indeed, the microtubule destabilizer Nocodazole promoted proinsulin secretion and cooperated with cytokines to further induce proinsulin secretion. To the best of our knowledge this is the first demonstration that cytokines and microtubule destabilization work together to promote proinsulin, as well as insulin, release.

One suggested mechanism for proinsulin secretion in T1D is that immature granules are released in which there is reduced expression of PCSK1 and CPE that process proinsulin maturation to insulin (21). Our finding that both proinsulin and insulin are released by cytokines is consistent with release of partially matured granules. Further, cytokines increased the interaction between proinsulin and the autoantigen ICA1(ICA69) that resides on the membranes of immature but not mature secretory granules (51). In contrast to the model in which there is a decrease in processing enzymes however, cytokines did not alter proinsulin interaction with PCSK1, and a recent proteomic study of human islets with the same cytokine treatment we used, did not find that cytokines altered PCSK1 and CPE protein levels (23). However, our results are consistent with the possibility that a subpopulation of misfolded proinsulin molecules deficient in anterograde trafficking may fail to reach a suitable compartment in which favorable interaction with PCSK1 or CPE may take place, even as another subpopulation of native proinsulin molecules may advance through the Golgi complex and interact favorably with processing enzymes.

The proinsulin AP-MS data also suggest an alternate microtubule dependent pathway leading to proinsulin secretion, through assembly and trafficking of stress granules (52). We found that cytokines strongly promoted proinsulin interactions with three highly conserved stress granule assembly factors; the diabetes GWAS factors Ataxin-2 and IGF2BP2, as well as the quintessential stress granule marker G3BP1 (34, 35, 53, 54). Stress granules are non-membranous cytoplasmic condensates formed via interactions among intrinsically disordered regions of proteins (55). They contain translationally halted transcripts as well as protein cargo that form spontaneously by liquid-liquid phase separation as part of the integrated stress response. How might proinsulin protein incorporation into stress granules result in its secretion? Notably: 1) stress granule protein components have been identified in extracellular vesicles (e.g. exosomes) from multiple cell types lines (56); 2) human islets secrete exosomes containing both proinsulin and insulin protein (57) and 3) IL-1β and IFN-γ treatment of human islets altered the composition of RNAs in exosomes, suggesting that changes in the protein composition may occur as well (58). Thus, the interactome data suggest that stress granules may also play a role in aberrant proinsulin trafficking/release.

The cytokine dependent proinsulin interactome data also provides clues related to T1D autoimmunity. In cytokine conditions, proinsulin interacted with TAP2 and Calnexin that together form an MHC intermediate complex during antigen presentation (47). Further, autoantigens

ICA69 and Chromogranin A bound proinsulin in cytokine conditions (39). Chromogranin A is a particularly intriguing proinsulin partner given that highly immunogenic peptides comprised of small regions of proinsulin and chromogranin A fused together were recently identified in T1D (59). It was suggested that molecular crowding in granules may contribute to the peptide-fusion phenomenon, and to the best of our knowledge the AP-MS data presented here provide the first demonstration that proinsulin and chromogranin A physically interact in human β-cells.

Taken together, the data reveal that even short-term exposure to inflammatory conditions reshapes the proinsulin interactome in ways that affect proinsulin folding, trafficking and secretion.

## Methods

### Human Islets

Islet preparations averaging 85-95% purity by DTZ and 95% viability by EB/FDA staining were procured from Prodo labs. The donors had no history of diabetes and had normal HbA_1c_. The islets were cultured for 48 hours in Prodo media of DMEM (105 mg/dL) with or without cytokines IL-1β (50U/ml) and IFN-γ (1000U/ml) for 48hrs. Nocodazole 5ug/mL was added to some cultures for 48 hours. PERK inhibitor GSK157 (2uM) was added to some cultures for 18-24 hours. SubAB and mSubAB (2.5 ug/mL) were added to some cultures for 6 hours.

### RNA-Seq

RNA was prepared from cytokine untreated and treated islets using RNeasy kit (Qiagen) and total RNA is Ribo depleted to remove rRNA from total RNA. The remaining non-rRNA is fragmented into small pieces using divalent cations under elevated temperature. Following fragmentation, the first strand cDNA was synthesized using random primers and followed by second strand synthesis using DNA Polymerase I. The cDNA is then ligated with index adapters for each sample followed by purification and then enriched with PCR to create the final library. The quality and quantity of the libraries were detected by Agilent Bioanalyzer and Kapa Biosystems qPCR. Multiplexed libraries are pooled and single-end 50-bp sequencing was performed on one flow-cell of an Illumina Next Gen Sequencing platform. For analysis, reads were mapped to human genome (version Homo_sapiens.GRCh37.75) and 87% overlap with exons was obtained. Then preferential association of 226 differentially expressed genes in treated vs non-treated condition with canonical pathways were mapped using Ingenuity Pathway Analysis.

### Proinsulin AP-MS

(Methods are described in detail in (11)). Briefly, human islets were lysed in 50mM Tris pH7.4, 150mM NaCl and 1% TX-100 with protease inhibitor cocktail (Thermo Fisher). Lysates were precleared with protein G agarose beads and immunoprecipitated with beads cross-linked to mouse IgG or proinsulin antibody (20G11) overnight at 4°C. A fraction of the beads were removed for protein elution to confirm successful IP by western blot and silver staining. The majority of the beads were subjected to denaturation, reduction and trypsin digestion followed by two-dimensional (2D) liquid chromatography–tandem mass spectrometry (LCMS/MS) analysis. The mass spectrometer was operated in positive data dependent acquisition mode with up to five MS2 spectra triggered per duty cycle. MS/MS spectra were searched against the Homo sapiens UniProt protein sequence database (January 2015 version) using MaxQuant (version 1.5.0.25). Enzyme was set to trypsin in a semi-specific mode and a maximum of two missed cleavages was allowed for searching. The target-decoy-based false discovery rate (FDR) filter for spectrum and protein identification was set to 1%. Intensity each identified peptide was subtracted from the control IP and Log 2Fold Change (Log2FC) was calculated. (Note that proinsulin was searched as insulin.) Label-free (LF) intensity was normalized by Loess method using Normalyzer. MSstats was used to calculate a confidence (*p* value) and fold change of proinsulin immunoprecipitation (IP)–to–control IgG IP intensity ratio for each protein. Human islet lysates were similarly immunoprecipitated for proinsulin for validation studies.

### Data Filtering

Datasets were filtered based on three criteria; Fold Change > 2, p< 0.05 and delta MS/MS counts > 10 in either untreated or cytokine treated condition. We then carried enriched ontology analyses on the interactors that were common to both groups or specific for Treated or Untreated showed enrichment for proteins belonging to “protein targeting” and “processing” functional groups (**Figure 2**).

### Western Blotting

Isolated islets were lysed in RIPA buffer (10mM Tris pH 7.4, 150mM NaCl, 0.1% SDS, 1% NP-40, 2mM EDTA) with protease and phosphatase inhibitors (Fisher Scientific) on ice for 30min and lysates were collected after centrifugation at 4°C for 15min at 12000g. Samples were prepared in Laemmli sample buffer without (non-reducing) or with (reducing) 5% beta-mercaptoethanol. After boiling for 5min, samples were analyzed by SDS-PAGE (4-12% Bis-Tris gel, (Bio-Rad Laboratories, Inc.) and transferred to nitrocellulose membranes (Bio-Rad Laboratories, Inc.). Primary antibodies used on human islets were as follows: proinsulin (20G11), BiP, Calnexin, phospho-EIF2α, EIF2α (EIFS1 gene) and Vinculin (Cell Signaling), GAPDH-HRP conjugate (Genetex), GBP5, IDO1, ERGIC1, Ataxin-2, ARFGAP2, and ERDJ3 (Proteintech), KIF2A and QSOX1 (Abcam) (see **Supplemental Table 1** for dilutions). For secondary antibodies, goat anti-mouse, goat anti-rabbit, were used in 1:5000 (Li-Cor, IRDye®-800CW or IRDye®- 680RD). After wash, membranes were imaged on Licor Odyssey CLX with ImageStudio software and for quantitation, the band intensities were analyzed by ImageJ.

### Immunohistochemistry

Human islets incubated for 48 hours +/-cytokines were harvested and fixed in 4% PFA. Paraffin embedding, sectioning, and slide preparations were done in the SBP Histopathology Core Facility. Sections were stained with primary antibodies (1:200) specific for: proinsulin (20G11, a kind gift of Dr. William Balch) and KIF2A (Abcam) and DAPI (Fisher Scientific). For secondary antibodies, Alexa Fluor 594 goat α-rabbit IgG, Alexa Fluor 488 goat α- mouse IgG were used (Invitrogen). Microscope images were taken at 200X.

### Griess Assay

Nitrite release into the media was tested using the Griess Reagent.

### ELISAs

IL6 (Abcam), proinsulin and insulin (Mercodia) were run according to manufacturer’s instructions.

## Acknowledgements

We thank Dr. William Balch for 20G11 antibody. This work was supported by NIH R24 DK110973-01 (PI-A, PA and RJK) and JDRF 2-SRA-2015-47-M-R, (PI-A, RJK and RJK) The project described was also supported by Helmsley Charitable Trust/ University of Miami Award Number 2018-T1D060 (PI-A). The content is solely the responsibility of the authors and does not necessarily represent the official views of the Helmsley Charitable Trust or University of Miami. We thank Alexandre Rosa Campos in the SBP proteomics core for helpful discussion and William Balch at Scripps Research Institute for 20G11 antibody.

