## Supplementary material for "Inflammatory cytokines rewire the proinsulin interaction network in human islets": Supp Material

##### Supplemental Figure Legends

**Supp Figure 1.** A) Western blot analysis protein expression in lysate and media; Vinculin, Indoleamine 2,3-Dioxygenase 1 (IDO1) and Guanylate Binding Protein 5 (GBP5) and proinsulin. Inset is second Western blot for proinsulin in media samples B) Intracellular proinsulin measured by ELISA and normalized to lysate GAPDH.

**Supp Figure 2.** A) Western blot and silver stain of proinsulin immunoprecipitations from human islets +/- cytokines. B) Total MS/MS counts from proinsulin AP/MS (blue) and IgG AP-MS of six human islets +/- cytokines. C) The total number of proteins identified by AP-MS in distinct islet samples labeled Exp. 6,7,10, 13,14,15. For each islet sample there are 4 conditions: Untreated/proinsulin IP, Untreated/IgG IP, Cytokine treated/proinsulin IP, and Cytokine treated/IgG IP. Note that biological samples 13, 14, 15 have two technical replicate MS runs each (R1 and R2).

**Supp Figure 3. AP-Western validations for selected Proinsulin Interactors identified by AP-MS.** Human islet lysates were immunoprecipitated with antibodies to proinsulin (or IgG), followed by Western blot for ERGIC1, Ataxin-2, ARFGAP2, ERDJ3, QSOX1 Calnexin and proinsulin.

**Supp Figure 4. Western blot for pEIF2$\alpha$ and total EIF2$\alpha$.** Human islets, untreated (CON) or cytokine treated (CYTO) were immunoblotted for EIF2$\alpha$ phosphorylated at ser51 (pEIF2 $\alpha$) and for total EIF2$\alpha$. MIN6 cells treated with Thapsigargin (Tg) were used as a positive control for pEIF2$\alpha$. GAPDH served as a loading control.

**Supp Figure 5. KIF2A versus Insulin expression in single human $\beta$-cells.** For single-cell RNA-Seq data obtained from Gene Expression Omnibus Repository (accession number GSE83139) the normalized reads for insulin (INS), glucagon (GCG) and KIF2A were log2 transformed. We defined $\beta$-cell identity as cells with INS log2 normalized reads >12 and with GCG log2 normalized reads <10. By these criteria, 99 cells were chosen for visualization of plotted INS values overlaid with KIF2A values.

### Supplemental figures

#### Inflammatory cytokines rewire the proinsulin interaction network in human islets

Duc Tran, Anita Pottekat, Kouta Lee, Megha Raghunathan, Salvatore Loguercio, Saiful Mir, Adrienne W. Paton, James C. Paton, Peter Arvan, Randal J. Kaufman, Pamela Itkin-Ansari

### Supplemental Figure 1

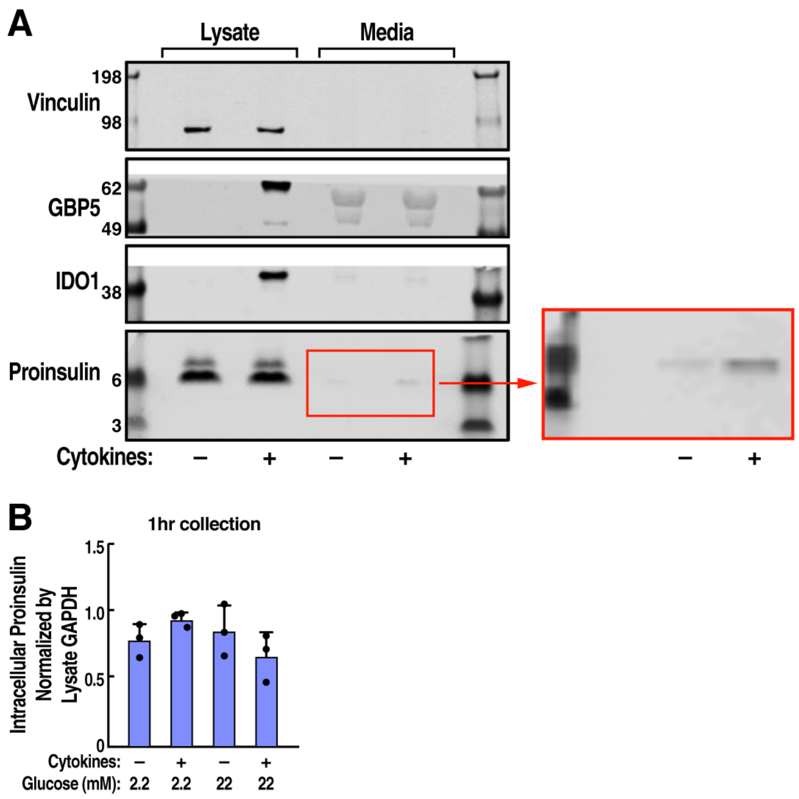

### Supplemental Figure 2

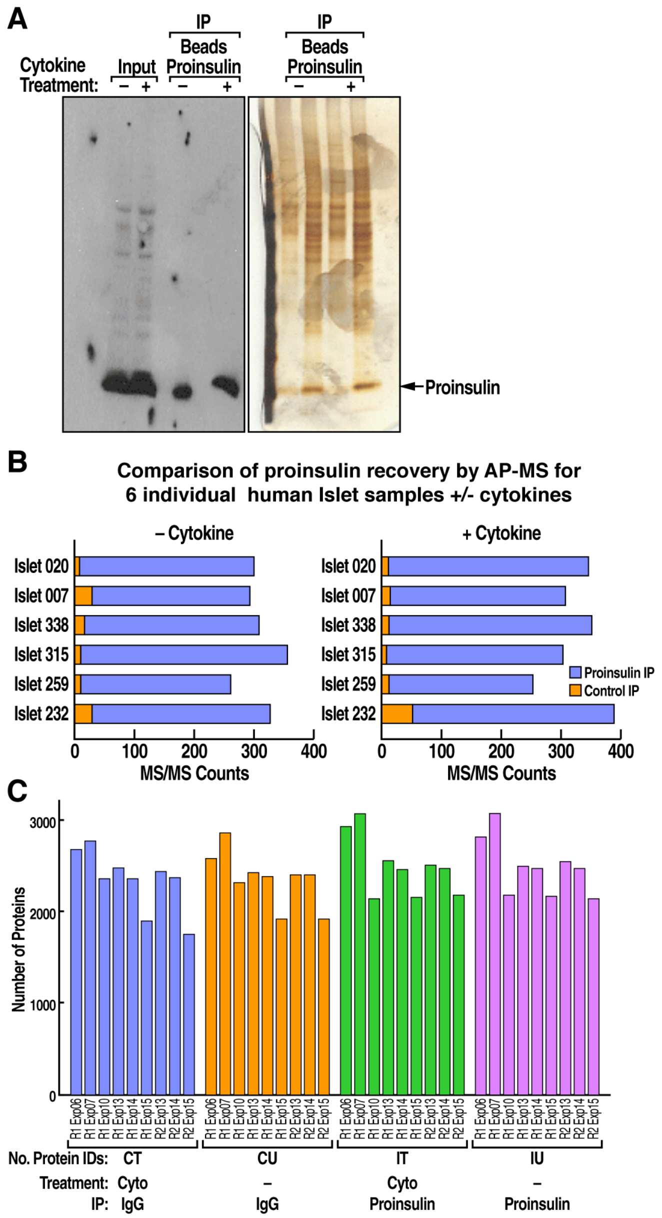

### Supplemental Figure 3

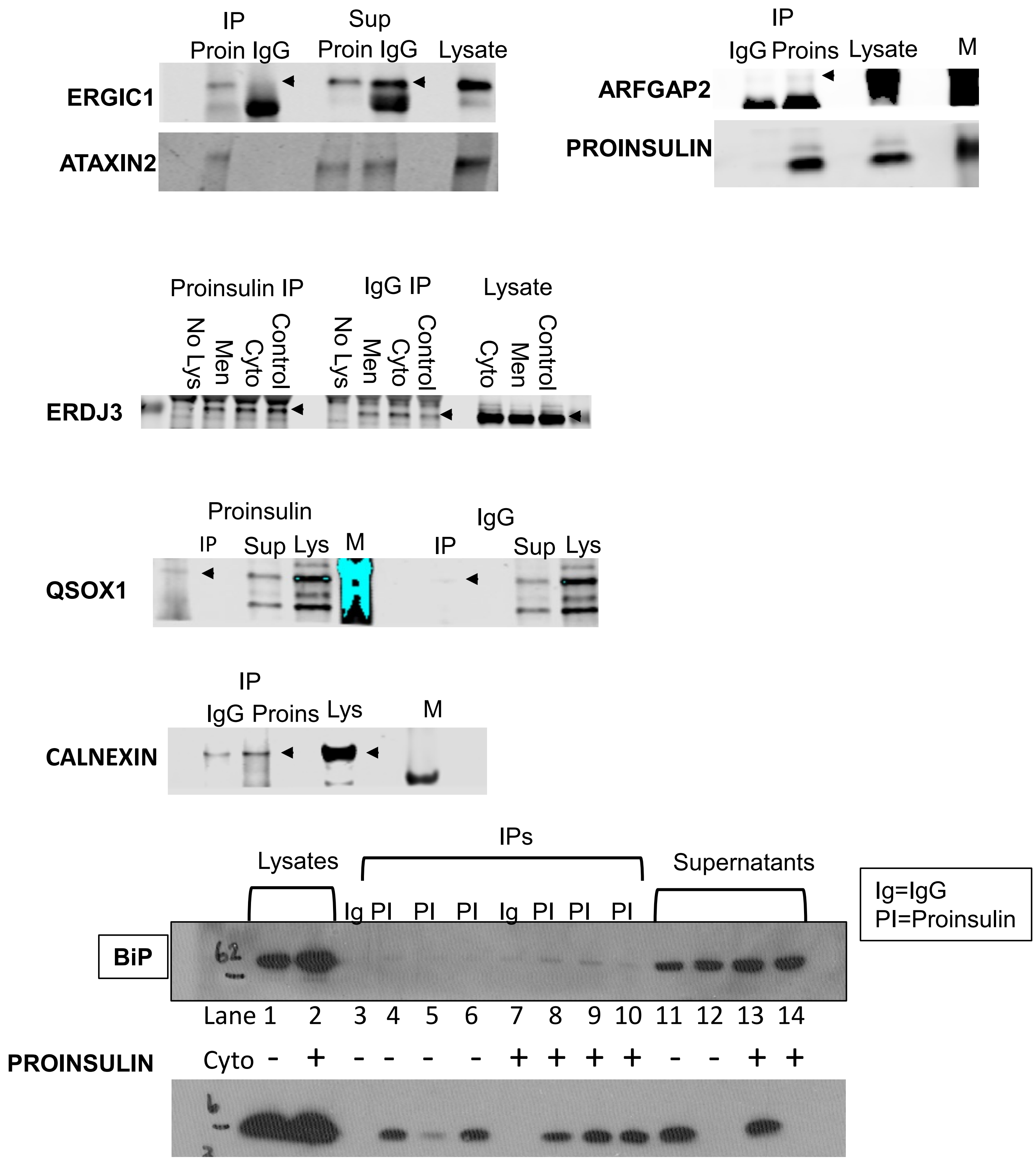

### Supplemental Figure 4

Cytokines do not induce EIF2$\alpha$ phosphorylation

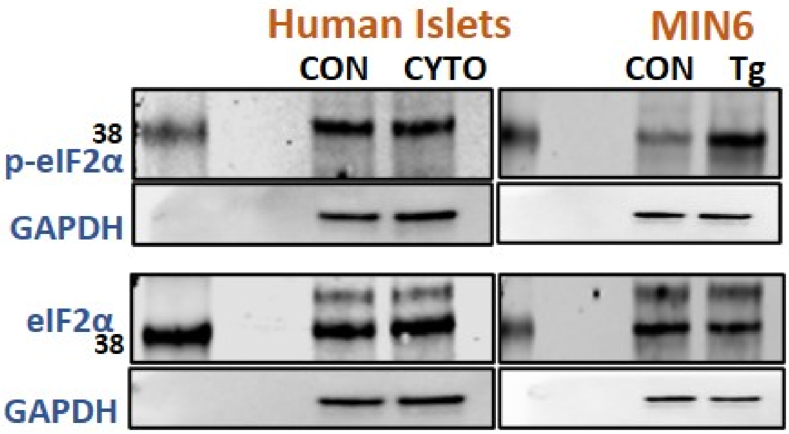

### Supplemental Figure 5

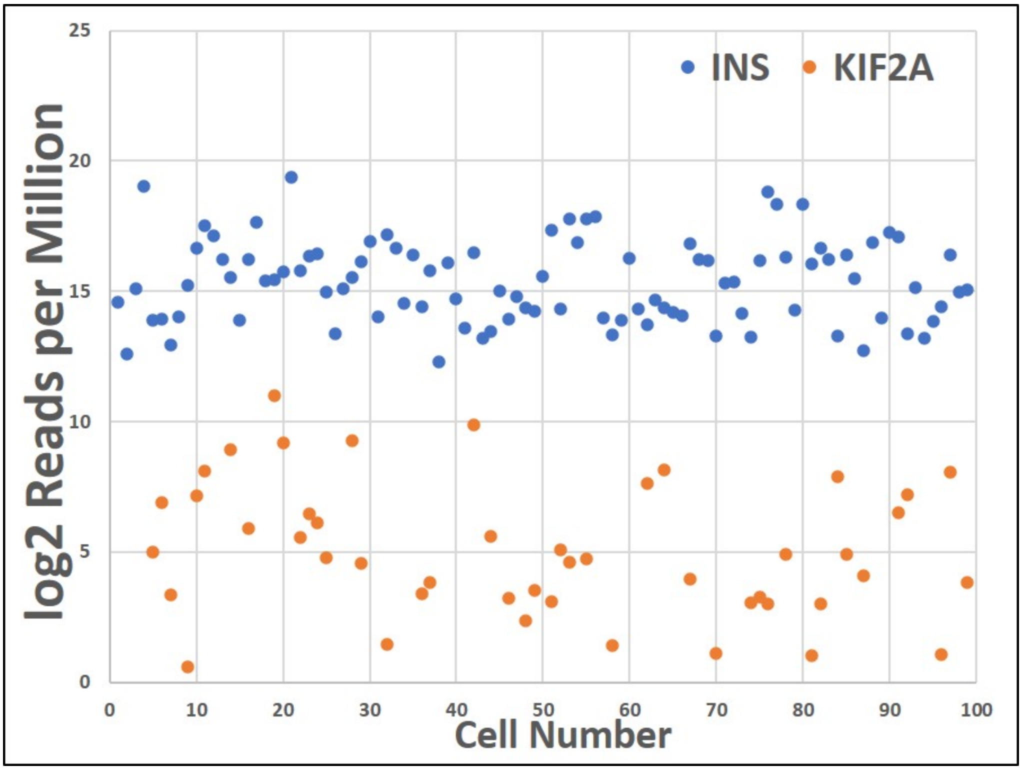

### Supplemental Table 1

Supplementary Table 1 - Primary antibodies

| Antibodies | Source | Product Number | Species | Dilution Factor |
|---|---|---|---|---|
| Anti-GBP5 | ProteinTech | 13220-1-AP | Rabbit | 1/1000 |
| Anti-IDO1 | ProteinTech | 13268-1-AP | Rabbit | 1/1000 |
| Anti-human proinsulin (20G11) | Dr William E. Balch's Lab | | Mouse | 1/1000 |
| Anti-human BiP | Cell Signaling | (C50B12) 3177 | Rabbit | 1/1000 |
| Anti-GAPDH-HRP | GeneTex | GTX627408-01 | Mouse | 1/100,000 |
| Anti-KIF2A | Abcam | ab197988 | Rabbit | 1/1000 for Western, 1/200 for IHC |
| Anti-vinculin | Cell Signaling | 4650S | Rabbit | 1/1000 |
| Anti-ERGIC1 | ProteinTech | 16108-1-AP | Rabbit | 1/1000 |
| Anti-ATAXIN2 | ProteinTech | 21776-1-AP | Rabbit | 1/5000 |
| Anti-ARFGAP2 | ProteinTech | 16519-1-AP | Rabbit | 1/600 |
| Anti-ERDJ3 | ProteinTech | 15484-1-AP | Rabbit | 1/1000 |
| Anti-QSOX1 | Abcam | ab235444 | Rabbit | 1/1000 |
| Anti-Calnexin | Cell Signaling | 2433S | Rabbit | 1/1000 |
| Anti-phospho-eIF2-alpha | Cell Signaling | 9722 | Rabbit | 1/1000 |
| Anti-eIF2-alpha | Cell Signaling | 9721 | Rabbit | 1/1000 |
